# PacBio library preparation using blunt-end adapter ligation produces significant artefactual fusion DNA sequences

**DOI:** 10.1101/245241

**Authors:** Paul Griffith, Castle Raley, David Sun, Yongmei Zhao, Zhonghe Sun, Monika Mehta, Bao Tran, Xiaolin Wu

## Abstract

Pacific Biosciences’ (PacBio) RS II sequencer, utilizing Single-Molecule, Real-Time (SMRT) technology, has revolutionized next-generation sequencing by providing an accurate long-read platform. PacBio single-molecule long reads have been used to delineate complex spliceoforms, detect mutations in highly homologous sequences, identify mRNA chimeras and chromosomal translocations, accurately haplotype phasing over multiple kilobase distances and aid in assembly of genomes with complex structural variation. The PacBio protocol for preparation of sequencing templates employs blunt-end hairpin adapter ligation, which enables a short turnaround time for sequence production. However, we have found a significant portion of sequencing yield contains chimeric reads resulting from blunt-end ligation of multiple template molecules to each other prior to adapter ligation. These artefactual fusion DNA sequences pose a major challenge to analysis and can lead to false-positive detection of fusion events. We assessed the frequency of artefactual fusion when using blunt-end adapter ligation and compared it to an alternative method using A/T overhang adapter ligation. The A/T overhang adapter ligation method showed a vast improvement in limiting artefactual fusion events and is now our recommended procedure for adapter ligation during PacBio library preparation.

## Introduction

Disease genomics and mRNA spliceoform analysis in cancer studies are benefitted by accurate long-read technologies which can be used to identify alternative splicing of mRNAs, fusion and translocation events, and to accurately sequence repetitive and unstable genes that are too complicated to be resolved by short-read technology (Haïg Djambazian, 2014; Rhoads & Au, 2015). Ligation of hairpin adapters onto both ends of the sequencing template molecules enables circular sequencing to produce multiple sequencing passes of each molecule which can be overlaid to build a highly accurate consensus sequence. Pacific Biosciences’ (PacBio) RSII system can produce accurate circular consensus sequence reads approaching 20 kb in length, with recently reported maximum polymerase read lengths of 92.7 kb (Nakano et al., 2017; Zhang et al., 2017). The ability to resolve diverse, complex structures which confound short-read technologies has created a preference over Illumina sequencing in specific applications that call for linked, long-range information. The PacBio library preparation protocol employs blunt-end ligation of the hairpin adapters, which raises a major concern that any two blunt-end DNA pieces may be ligated together to produce artefactual fusion molecules and concatamers, prior to adapter ligation. The protocol calls for an excess of blunt-end adapter compared to template molecules, which is thought to prevent template concatamer formation. However, we have found that this issue needs to be addressed carefully.

Use of blunt-end adapter ligation during library preparation enables users to complete library construction and begin sequencing on the PacBio RS II in less than 24 hours (PacificBiosciences, 2015). This short sample-to-sequence time is made possible by a blunt-end adapter ligation reaction that can be completed in 15-60 minutes. Early PacBio library construction protocols utilized an overnight A/T overhang adapter ligation reaction for insert fragments less than 500 bp but these have since been discontinued from marketing and reference literature in favor of the faster blunt-end adapter ligation method. Concern related to blunt-end ligation is focused on the potential for concatamer formation comparable to that seen with Ion Torrent ligation-based library preparation protocols (Gorbacheva, Quispe-Tintaya, Popov, Vijg, & Maslov, 2015). Recent studies of mRNA isoform diversity have reignited this concern regarding the PacBio platform, where detailed analysis of long reads demonstrated that 33% of all analyzed reads were concatamers of shorter fragments (Karlsson & Linnarsson, 2017). Furthermore, these concatamers existed primarily as three or more individual fragments within a single Circular Consensus Sequence (CCS) read (Karlsson & Linnarsson, 2017).

Our study originally focused on NCF1, a gene involved in 22% of all CGD patients and the predominant cause of the autosomal recessive form of the disease (Roos et al., 2006). This locus presents a complicated sequencing situation with two pseudogenes (ΨNCF1B and ΨNCF1C) and a single functional copy (NCF1A) which share >98% homology (Chanock et al., 2000). Assembly of short-read sequences cannot properly differentiate NCF1A mutations or SNPs from ΨNCF1B or ΨNCF1C, precluding mutation identification. Use of single-molecule sequencing provides long, accurate reads that enable identification and linkage of novel mutations with NCF1A associated SNPs. To this end, evaluation of the accuracy of library preparation for NCF1 is a critical point in the development of an assay for mutational analysis.

To evaluate the potential for concatamer formation prior to our own study, we produced 1-1.5 kb PCR fragments amplified from the pBR322 plasmid. The PCR products were split in half for PacBio library preparation, applying blunt-end adapter ligation to one half and A/T overhang adapter ligation to the second half. The implications of analyzing concatamer formation across these size ranges may be significant across multiple PacBio applications, and provide insight into the risk of concatamer formation in our studies.

## Methods

**PCR of pBR322 fragments for PacBio sequencing** was performed for size specific amplification and M13 tagging. PCR utilized 12.5μL 2X Q5 Hot Start master mix (New England Biolabs) 1.25μL forward primer (10 μM), 1.25μL reverse primer (10 μM), 1μL pBR322 vector (New England Biolabs) and 14μL H_2_O. Primer combinations can be found in Table 1. Following confirmation of fragment size via 1.2% Flash-Gel (Lonza), samples were purified using Agencourt AMPure XP beads (Beckman Coulter) and analyzed on a Bioanalyzer DNA 1000 or DNA 7500 chip (Agilent), as appropriate.

**Table 1.**
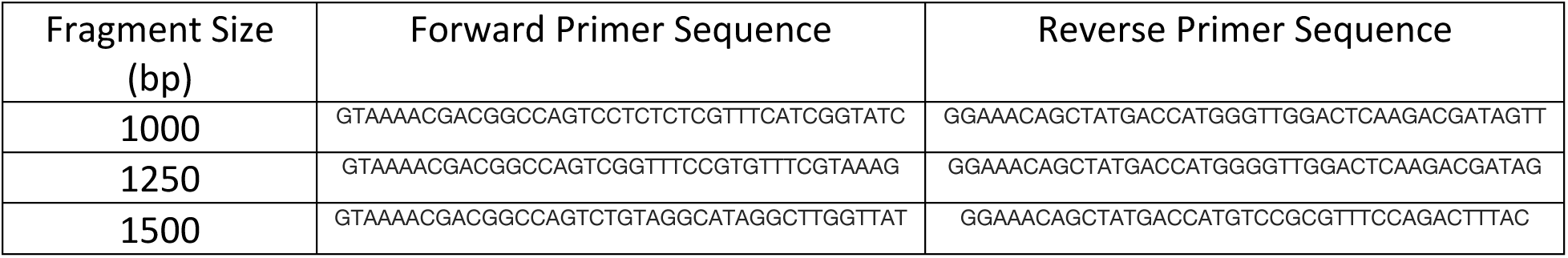
Primer sequences utilized for amplification of pBR322 for PacBio sequencing.

**PCR of NCF1 for PacBio Sequencing** was done using two rounds of PCR, in a manner similar to pBR322 amplification and tagging. The first round used a 20μL mix of 2μL 10X Platinum Taq buffer (Invitrogen), 0.6μL 50mM MgCl_2_, 0.4μL 10mM dNTP mix, 0.4μL 10μM forward primer (10 μM) (AGTAAAACGACGGCCAGTGAGAAGCGCTTCGTACCCG), 0.4μL of 10μM reverse primer (10 μM) (GGAAACAGCTATGACCATGCGTCCAGACGCCAGGCTCTATA), 0.8μL Platinum Taq (Invitrogen) 2μL 1^st^-strand cDNA and H_2_O up to 20μL to amplify NCF1 with M13 tags. The second cycle added sample-specific barcodes to the M13-tagged NCF1 PCR product using 2μL 10X Platinum Taq buffer (Invitrogen), 0.6μL 50 mM MgCl_2_, 0.4μL 10 mM dNTP mix, 0.4μL of 5μM forward primer (10 μM), 0.4μL of 5μM reverse primer (10 μM), 0.8μL Platinum Taq (Invitrogen), 1μL 1^st^-round PCR product and H_2_O up to 20μL. PCR products generated contained two 8 bp barcode regions flanking NCF1 (Figure 1). PCR products were pooled in an equimolar solution prior to library preparation.

**Figure 1.**
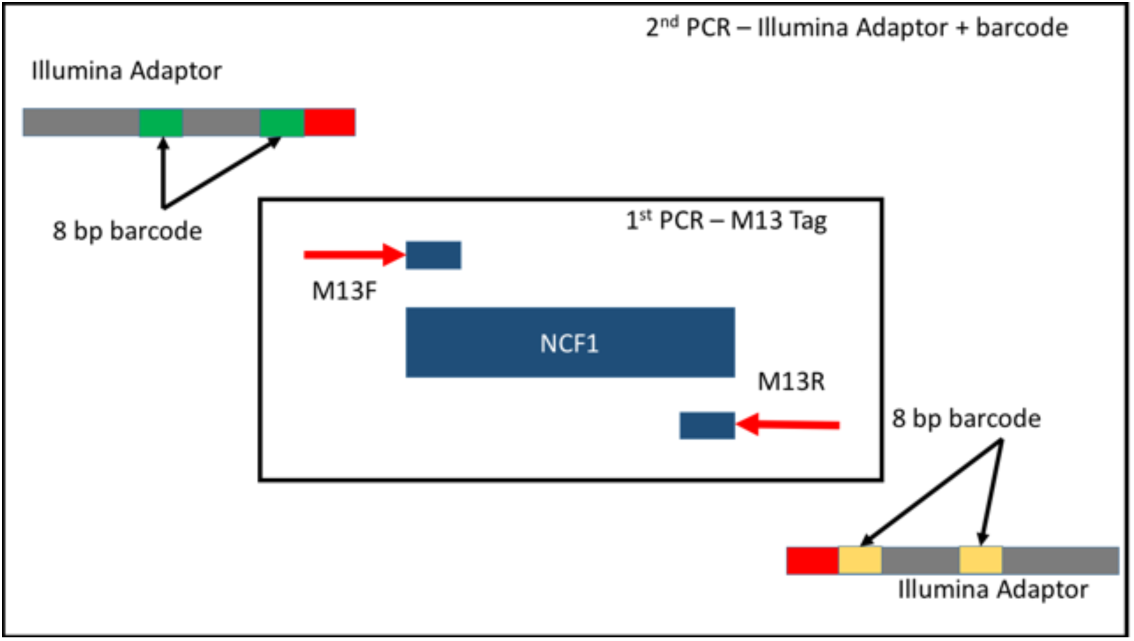
Illustration of barcode addition to NCF1 cDNA. Sample cDNA was tagged with M13 sequences (red) in the first PCR reaction. A second PCR reaction was performed to add Illumina adapters (grey) with two 8 bp barcodes (green and yellow) on the 5’ and 3’ ends of the cDNA.

**PacBio Library Preparation** of NCF1 PCR samples was done in accordance with PacBio standards as laid out in the PacBio user’s manual. 6 libraries were prepared prior to sequencing. Two libraries, one utilizing PacBio’s standard blunt-end ligation protocol (PN 001-143-835-08) and the other utilizing PacBio’s A/T overhang ligation protocol (PN 001-214-501-03), were generated separately for 1 kb fragments and 1.5 kb fragments prior to equimolar pooling. The damage repair step prior to end repair was included in the A/T overhang ligation protocol. This step was not included in PacBio’s original A/T overhang ligation protocol. The two pooled libraries containing either blunt-end or A/T overhang ligation products were then sequenced in two SMRT cells, respectively. Two additional libraries were prepared using blunt-end and A/T overhang ligation preparations by pooling 1 kb, 1.2 kb and 1.5 kb fragments in equimolar ratios prior to library preparation and subsequent sequencing in two SMRT cells. Blunt-end ligations were performed as per the standard PacBio protocol, 001-143-835-08. A-tailing of PCR fragments was done using Klenow (exo -) fragment for 60 minutes prior to A/T overhang ligation following the PacBio protocol 001-214-501-03. A/T overhang adapter ligation was incubated at RT overnight, prior to ligase inactivation.

**Sequence analysis of PacBio results** utilized CCS reads generated through sequencing, and classified complete, full-length reads when the aforementioned M13 sequences were recognized at both ends of the sequence of interest. To prevent misidentification of M13 primer sequences due to potential sequencing errors, the hamming distance between the M13 primer sequences allowed 3 mismatches and no ambiguity as to which M13 primers sequences a given read belonged to. Following filtering of incomplete sequences that did not possess at least two M13 tags, sequences were analyzed for total number of independent M13 pairs. The resulting CCS reads utilized for concatamer analysis was lower than the original CCS read count due to this filtering. Concatamers were determined by visualization of multiple pairs of M13 tags from separate samples within a single CCS read.

## Results

### PCR-generated pBR322 fragments produce concatamers during standard PacBio library preparation, prior to loading onto PacBio RSII SMRT cells

PCR-generated pBR322 fragments of 1 kb, 1.2 kb and 1.5 kb served as estimators of library preparation concatamer generation for short, consistent length fragments. Following library preparation, fragments were analyzed via Agilent Bioanalyzer as a quality control step to confirm proper preparation, concentration and sizing. Bioanalyzer results depicted concatamer formation, resulting in bands double and triple the expected sizes of the pBR322 inserts. Both blunt-end ligation-based library preparations demonstrated noticeable peaks at 2x and 3x the expected insert sizes. These additional peaks were notably absent in libraries that were prepared using A/T overhang ligation (Figure 2). This initial assessment suggests that the formation of concatamers occurs in a blunt-end ligation specific manner prior to sequencing, with 7.06% of the total DNA in 1 kb/1.5 kb independently prepared (subsequently pooled) libraries representing concatamers. This percentage decreases to 6.50% of total library DNA when 1 kb, 1.2 kb and 1.5 kb fragments are pooled prior to blunt-end library preparation.

**Figure 2.**
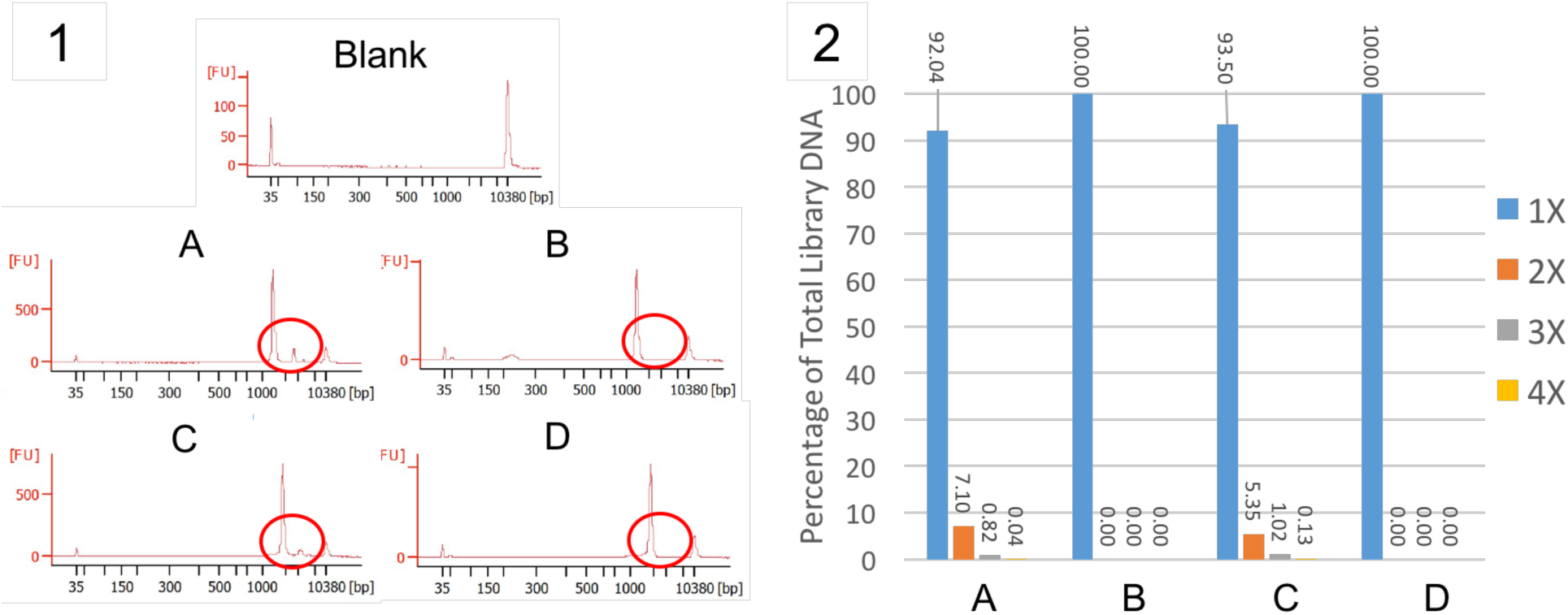
Analysis of concatamer formation during library preparation of 1 kb and 1.5 kb fragments prepared using blunt-end (A) or A/T overhang (B) ligation and pooled libraries of 1 kb, 1.2 kb and 1.5 kb fragments prepared using blunt-end (C) or A/T overhang (D) ligation. Concatamers are readily visible in blunt-end ligation (A, C) and absent in A/T overhang ligation (B, D) preparations (red circle) via Bioanalyzer High Sensitivity chip following library preparation (1). Concatamers represent a measurable percentage of the total library DNA (2) when using blunt-end library preparation (A, C), while they are not detectable via high sensitivity Bioanalyzer chip when using A/T overhang ligation library preparation (B, D).

### PCR amplicons of pBR322 of 1 kb and 1.5 kb lengths with separately prepared sequencing libraries see a decrease in concatamers present in CCS reads when using A/T overhang adapter ligation

CCS reads with more than one pair of M13 tags were counted and assessed for the total number of inserts present between PacBio adapters resulting from concatamerization of the inserts. The SMRT cell for the blunt-end ligation library generated 32,477 CCS reads containing at least one pair of M13 tags, 957 of which contained more than two M13 pairs. These reads represented the concatamers formed using blunt-end adapter ligation, equating to 2.85% of the total reads (Figure 3A). Conversely, the SMRT cell for the A/T overhang ligation library generated 33,144 CCS reads containing at least one pair of M13 tags, 26 of which contained more than two M13 pairs, equating to only 0.08% of the total CCS reads being concatamers (Figure 3B). A/T overhang ligation during library preparation provided a 97.19% reduction in total concatamer formation when libraries were generated with a single length fragment of either 1 kb or 1.5 kb prior to sequencing.

**Figure 3.**
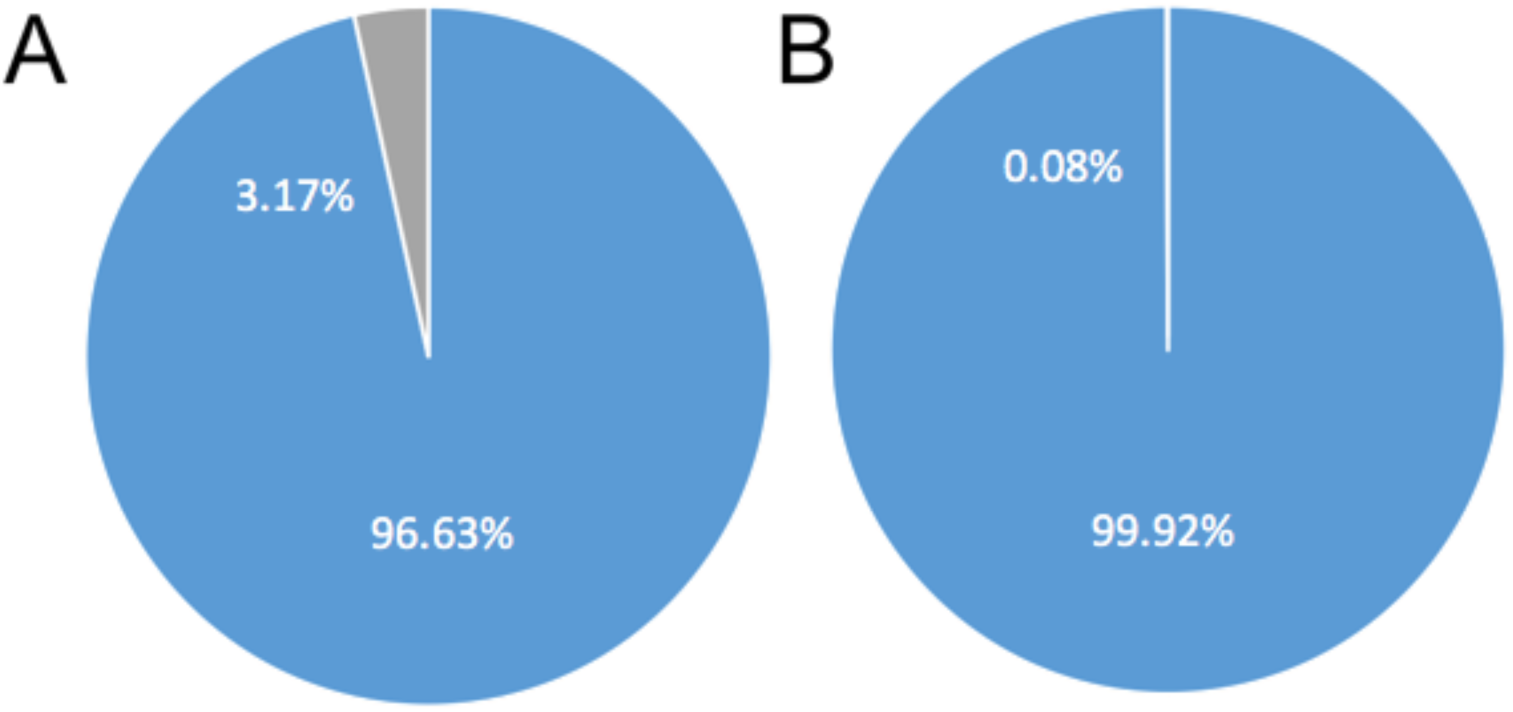
Percentage of CCS reads produced from PacBio RS II sequencing for 1 kb and 1.5 kb fragments using either blunt-end or A/T overhang ligated library preparation that contain concatamers (gray) vs. single inserts (blue). Following sequencing via PacBio RS II, CCS reads were analyzed for the inclusion of multiple tagged fragments within a single read, denoting the presence of concatamers. Libraries prepared using blunt-end ligation (A) produced considerably more CCS reads with concatamers than those prepared with A/T overhang ligation (B).

### Sequencing libraries from PCR fragments of pBR322 with lengths of 1 kb, 1.2 kb and 1.5 kb pooled and prepared using either blunt-end or A/T overhang adapter ligation demonstrated measurable differences in concatamer formation

Assessment of the total number of CCS reads from inserts generated by concatamers was done by studying pairs of M13 tags which were placed at the outside of each original PCR fragment. One SMRT cell for the blunt-end ligation generated pooled library resulted in 27,662 CCS reads with at least one set of tags, with 1,790 containing more than one pair of M13 tags, representing 6.05% of the total CCS reads (Figure 4A). The SMRT cell for the pooled library generated via A/T overhang ligation resulted in 26,893 CCS reads with at least one pair of M13 tags, 29 of which contained more than one pair, representing only 0.11% of the total CCS reads (Figure 4B). For pooled libraries, using A/T overhang ligation instead of blunt-end ligation reduced the incidence of concatamer formation by 98.18%.

**Figure 4.**
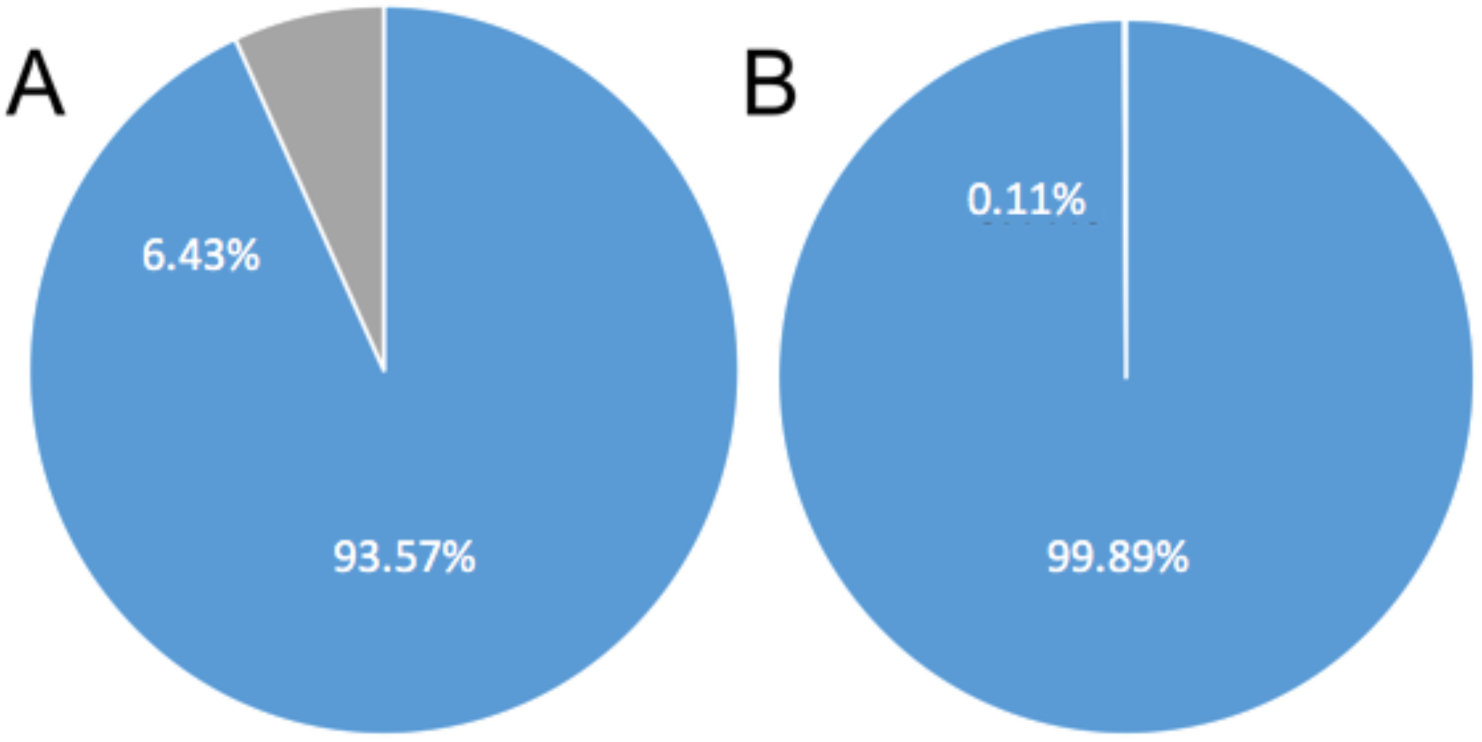
Percentage of CCS reads produced from PacBio RS II sequencing for pooled library preparations of 1 kb, 1.2 kb and 1.5 kb fragments using either blunt-end or A/T overhang ligated library preparation that contain concatamers (gray) vs. single inserts (blue). Following sequencing via PacBio RS II, CCS reads were analyzed for the inclusion of multiple tagged fragments within a single read, denoting the presence of concatamers. Libraries prepared using blunt-end ligation (A) produced considerably more CCS reads with concatamers than those prepared using A/T overhang ligation (B).

### Blunt-end and A/T overhang adapter ligation produce drastically different numbers of concatamers using PacBio sequencing of NCF1

CCS reads produced using blunt-end ligation (17,204 reads) and A/T overhang ligation (27,131 reads) were analyzed for the presence of two M13 pairs to classify reads as full-length. In the library prepared using blunt-end ligation, 24.4% of the total CCS reads that were classified as full-length were recognized as concatamers, as determined by the presence of multiple barcode pairs in a single CCS read (Figure 5A). By stark contrast, the A/T overhang ligation prepared library was found to contain concatamers representing only 0.04% of the total CCS reads. This represents a total reduction of 99.84% in concatamer formation (Figure 5B). Further analysis of the concatamers produced by both blunt-end and A/T overhang adapter ligation revealed that, in addition to concatamers with two unrelated sample sequences, blunt-end ligation preparations produced concatamers with up to six separate barcoded sequences in a single CCS read, while A/T overhang ligation preparations produced only 13 concatamers with two separate barcoded sequences present (Table 2).

**Figure 5.**
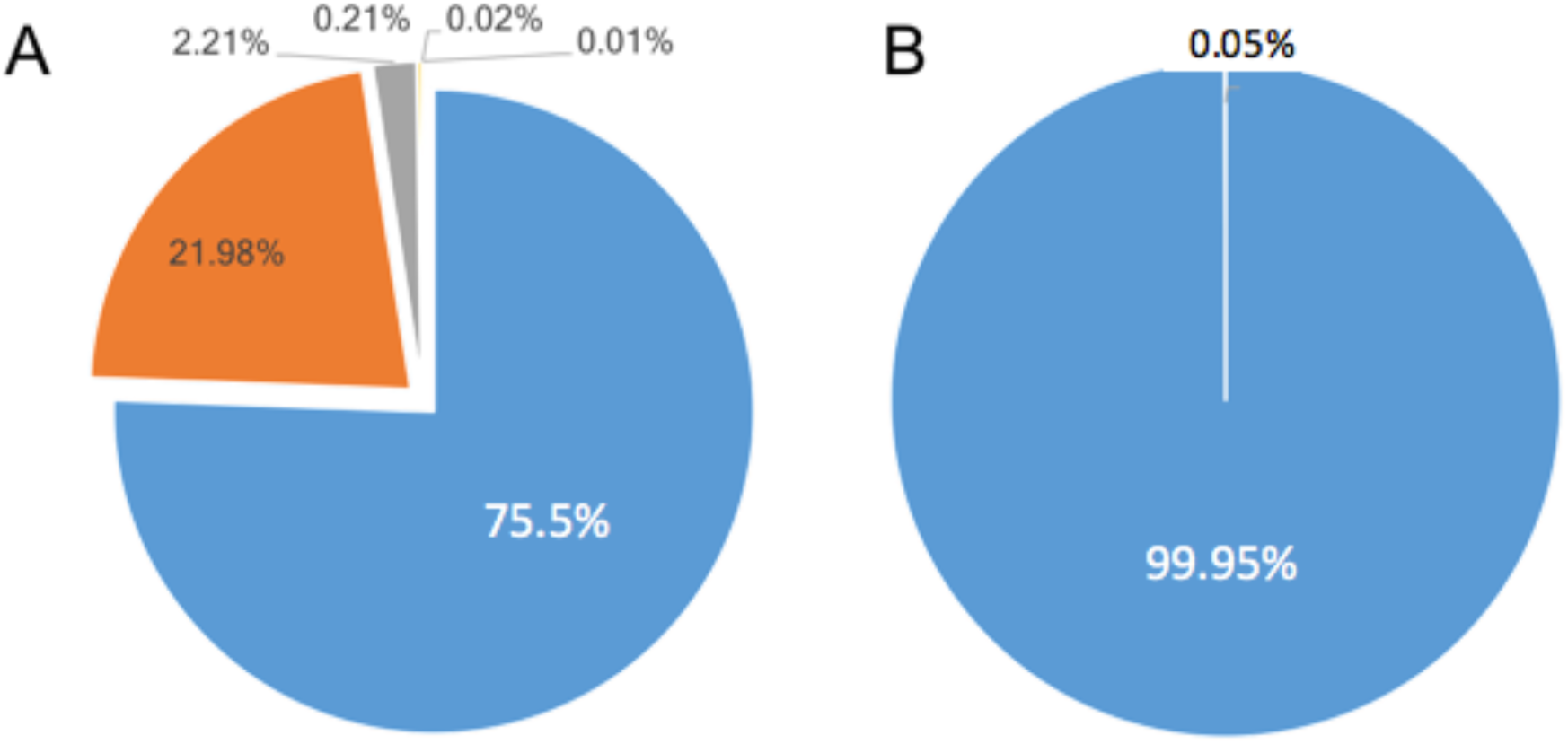
Concatamers formed as a proportion of total CCS reads using blunt-end or A/T overhang ligation strategies for NCF1. Figure 4A presents the proportion of CCS reads which were determined to be concatamers generated by using the standard PacBio blunt-end ligation preparation with 2 inserts (orange), 3 inserts (grey) and additional percentages listed across the top representing (L to R) 3 insert concatamers, 4 insert concatamers, 5 insert concatamers and 6 insert concatamers. Figure 4B presents the results from A/T overhang adapter ligation.

**Table 2.** Total number of NCF1 CCS reads containing one or more complete barcoded sequences generated from independent library preparations of 1 kb and 1.5 kb fragments, pooled 1 kb, 1.2 kb and 1.5 kb libraries and NCF1 libraries.

| Unique M13-Tagged Within a Single CCS Read | 1 kb and 1.5 kb Fragments: Blunt-End Ligation Library Preparation | 1 kb and 1.5 kb Fragments: TA-Ligation Library Preparation | Pooled Fragments: Blunt-End Ligation Library Preparation | Pooled Fragments: TA-Ligation Library Preparation | NCF1 Blunt-End Ligation Library Preparation | NCF1 TA Ligation Library Preparation |
| --- | --- | --- | --- | --- | --- | --- |
| 1 | 32477 (96.63%) | 33144 (99.92%) | 27662 (93.57%) | 26893 (99.89%) | 17204 (75.5%) | 27131 (99.95%) |
| 2 | 957 (2.85%) | 26 (0.08%) | 1790 (6.05%) | 29 (0.11%) | 5005 (22.0%) | 13 (0.05%) |
| 3 | 94 (0.28%) | 0 | 92 (0.31%) | 0 | 503 (2.2%) | 0 |
| 4 | 12(0.04%) | 0 | 14 (0.05%) | 0 | 48 (0.2%) | 0 |
| 5 | 1(<0.01%) | 0 | 5 (0.02%) | 0 | 4 (0.01%) | 0 |
| 6 | 0 | 0 | 0 | 0 | 2 (0.005%) | 0 |

## Discussion

Assembly of products sequenced on the PacBio RS II platform rely on accurate interpretation of the reads of insert produced by circular consensus sequencing. Our results call into question the use of blunt-end adapter ligation for PacBio library preparation due to the potential for widespread concatamer formation. Although many analysis pipelines employ filters to remove concatamers and erroneous sequences in the post-sequencing evaluation of CCS reads, these pipelines often require a reference. The presence of concatamers may introduce confounding information when attempting assembly of novel genomes, construction of alternative spliceoforms, or resolving genomic rearrangements. Therefore, for analysis of unknown sequences lacking a definitive reference, the current method of blunt-end library preparation should be evaluated against A/T overhang library preparation. Our data suggests that the risk of concatamer formation is a genuine concern for analysis of fragments less than 1.5 kb, and more importantly, this complication is readily resolved using A/T overhang adapter ligation.

Fidelity during library preparation is key for sequencing on any platform. Both library preparation methods generated ligation products within single pairs of PacBio adapters, however, use of A/T overhang adapter ligation reduced the incidence of concatamer appearance in CCS reads by 98.65% ± 1.09%. This reduction is most apparent in the disappearance of concatamers of more than two PCR fragments per CCS read, which we found to be commonplace in blunt-end ligated libraries, but not present in A/T overhang ligated libraries. We suspect that the total number of concatamers formed during library preparation, including those concatamers containing more than two fragments, are under-represented in the CCS reads due to the preference for shorter fragments to load in the SMRT cell.

Expanding on our controlled experiment, looking at an uncontrolled, sample-derived PCR amplicon from NCF1 illustrates the potential for significant variation in concatamer formation depending on currently unforeseen sequence-based differences. Although our controlled experiments with pBR322 fragments demonstrated concatamer formation, the total percentage of CCS reads containing concatamers is below 7%. When expanding this analysis to NCF1 we see an increase in concatamer formation to approximately 24% of the total reads. As with pBR322, the ability to reduce the total number of concatamers by over 99% is available by using A/T overhang adapter ligation. We have additional anecdotal evidence that other sample-derived gene preparations also resulted in high numbers of concatamers due to the use of blunt-end adapter ligation, but these have not been studied under controlled settings as we have done above. We propose that laboratories that are using the RS II system for the purposes of discovery of genomic rearrangements or Iso-seq analysis evaluate the use of A/T overhang adapter ligation in future library preparations to reduce the significant risk of concatamer formation.

## Acknowledgements

This project has been funded in whole or in part with Federal funds from the National Cancer Institute, National Institutes of Health, under Contract No. HHSN261200800001E. The content of this publication does not necessarily reflect the views or policies of the Department of Health and Human Services, nor does mention of trade names, commercial products, or organizations imply endorsement by the USA Government.

We would also like to acknowledge Pacific Biosciences of California for providing their previously marketed A/T overhang ligation adapters for this analysis.

